# Characterisation and automated quantification of induced seizure-related behaviours in *Xenopus laevis* tadpoles

**DOI:** 10.1101/2022.11.18.517140

**Authors:** Sandesh Panthi, Phoebe A. Chapman, Paul Szyszka, Caroline W. Beck

**Affiliations:** Department of Zoology, University of Otago, Dunedin, New Zealand, 9054; Brain Health Research Centre, University of Otago, Dunedin, New Zealand; Genetics Otago, University of Otago, Dunedin, New Zealand

**Keywords:** Epilepsy, seizure behaviour, African Clawed Frog, *Xenopus laevis*, TopScan

## Abstract

Epilepsy, a clinical diagnosis characterized by paroxysmal episodes known as seizures, affects 1% of people worldwide. Safe and patient-specific treatment is vital and can be achieved by the development of rapid pre-clinical models of for identified epilepsy genes. Epilepsy can result from either brain injury or gene mutations, and can also be induced chemically. *Xenopus laevis* tadpoles could be a useful model for confirmation of variants of unknown significance found in epilepsy patients, and for drug re-purposing screens that could eventually lead to benefits for patients. Here, we characterise and quantify seizure-related behaviours in *X. laevis* tadpoles arrayed in 24-well plates. To provoke acute seizure behaviours, tadpoles were chemically induced with either pentylenetetrazole (PTZ) or 4-aminopyridine (4-AP). To test the capacity to adapt this method for drug testing, we also exposed induced tadpoles to the anti-seizure drug valproate (VPA). Four induced seizure-like behaviours were described and manually quantified, and two of these (darting, circling) could be accurately detected automatically, using the video analysis software TopScan. Additionally, we recorded swimming trajectories and mean swimming velocity. Automatic detection showed that either PTZ or 4-AP induced darting behaviour and increased mean swimming velocity compared to untreated controls. Both parameters were significantly reduced in the presence of VPA. In particular, darting behaviour was a shown to be a sensitive measure of epileptic seizure activity. While we could not automatically detect the full range of seizure behaviours, this method shows promise for future studies, since *X. laevis* is a well-characterised and genetically tractable model organism.

## Introduction

Epilepsy is a common neurological condition characterised by recurrent seizures, which are transient episodes of abnormal excessive brain activity (Fisher *et al*. 2005). Currently, 50 million people are estimated to have epilepsy and five million people are diagnosed with epilepsy each year (Fiest *et al*. 2017; Fisher *et al*. 2014). Epilepsy can be classed as either acquired (e.g. following injury or trauma) or genetic in origin. Genetic epilepsy incidence peaks within the first year of life (Camfield & Camfield 2015). Children under 5 years of age have the highest age-specific incidences (∼60 per 100,000) (Symonds *et al*. 2021). Developmental and epileptic encephalopathies (DEE) are the most severe forms of childhood epilepsy with frequent severe epileptiform activity and developmental slowing (McTague *et al*. 2016). DEE increases the risk of premature mortality and patients who survive have significant lifelong disabilities (Palmer *et al*. 2021). Patients are normally treated with one or more anti-seizure medications (ASM) which aim to reduce the frequency of seizures. However, ASM are not always sufficient and at best they only suppress seizure activity, with the underlying developmental defect untreated. Safe and effective patient-specific approaches are required for the treatment of DEE, ideally targeting the specific molecular pathways that are affected by specific gene mutations (Loscher 2020).

Recent advances in molecular genetics have led to the discovery of more than 140 epilepsy-associated genes or loci (Perucca *et al*. 2020; Epi4K *et al*. 2013). Within each defective gene there are unique pathogenic variants, and phenotypic variation is common (Begemann *et al*. 2021; Indelicato & Boesch 2021; Sadleir *et al*. 2017). This combination of a large number of genes and variants within genes means developing pre-clinical seizure models is a huge challenge in terms of time, cost and efficient results. Being able to develop models that can be used to both validate potential variants as pathogenic and quickly assess conventional or repurposed drugs for their ability to reduce seizure frequency will help in developing drugs against genetic epilepsy.

To date, various animal models of epilepsy/epileptic seizures have been established and characterized (Grone & Baraban 2015; Loscher 2017). Genetic and pharmacological rodent models of epilepsy have greatly contributed on the genetics of epilepsy and strongly improved our understanding of the pathophysiology of this neurological disorder (Grone & Baraban 2015). However, every model has its benefits and its limitations (Kandratavicius *et al*. 2014) and functional studies of the growing number of epilepsy-causing genes and variants is failing to keep pace with gene discovery. Recent approaches take advantage of the smaller, more fecund aquatic vertebrate models *Danio rerio* (Zebrafish) and *Xenopus laevis* (South African clawed frog), both of which are already well-established models for behavioural neuroscience and are additionally amenable to genetic manipulation (Hortopan *et al*. 2010; Pratt & Khakhalin 2013; Adamson *et al*. 2018; Vaz *et al*. 2019; Willsey *et al*. 2022).

*Xenopus* are prolific breeders, and their brain development is very well characterised (Exner & Willsey 2021). The development from eggs to pre-feeding, transparent tadpoles with well-developed brains takes place in a week and can be controlled by manipulating temperature (Nieuwkoop & Faber 1994). The *X. laevis* genome has been sequenced and is well annotated (Session *et al*. 2016) and the Xenbase database supports research into genetic disease (Nenni *et al*. 2019). Further, *Xenopus* are highly amenable to manipulation such as CRISPR/Cas9 editing (Horb *et al*. 2019; Naert *et al*. 2020) or expression of exogenous proteins via mRNA injections (Gurdon *et al*. 1974). These unique properties make *X. laevis* tadpoles a viable model organism for the functional study of genetic epilepsy and epileptic seizures, as well as for pre-clinical drug testing.

The use of *Xenopus* tadpoles to model acute seizures (induced by chemicals) has been previously described, and a series of stereotypical behaviours were classified (Bell *et al*. 2011; Hewapathirane *et al*. 2008). The most severe behaviour involves a strong contraction of trunk muscles on one side of the tadpole such that its head and tail connect briefly. These movements are known as C-shaped contractions due to the characteristic extreme posture (Hewapathirane *et al*. 2008). Recently, CRISPR/Cas9-mediated knockdown in *Xenopus* tadpoles was used to confirm two *de novo* human NEUROD2 variants as causative for DEE-75 (Sega *et al*. 2019). Importantly, Sega and colleagues described both abnormal swimming and the spontaneous occurrence of C-shaped contractions (CSC) as seizure-related behaviour, showing that chronic epilepsy behaviours can also be phenotyped in this model (Sega *et al*. 2019). However, automated detection of these behaviours, which may be important if these models are to be useful in closing the gap between diagnosis and treatments, has not yet been described. Here, we evaluate the potential for a popular software for mammalian behaviour analysis, TopScan, to detect seizure-like behaviours in *X. laevis* tadpoles. Two acute seizure-inducing drugs were applied to tadpoles arrayed in 24-well plates in the presence or absence of anti-seizure drug valproic acid (VPA), and the resulting behaviours both manually and automatically tallied. While we could not reliably automatically detect the C-shaped contractions, other seizure-associated behavioural parameters could be automatically detected. Automatic behaviour tracking therefore adds to the toolkit that makes *X. laevis* tadpoles a promising model for rapid phenotyping of models of chronic epilepsy in future.

## 2 Materials and methods

### 2.1 Husbandry and production of *Xenopus laevis* eggs and embryos

*Xenopus laevis* adult males and females are maintained in standard aquarium conditions with 12/12h light dark cycles at 18 ºC in carbon filtered tap water at the Zoology Department, University of Otago. The colony has been closed since its establishment in 2004 and is now in the F2 generation. Adult female *X. laevis* (55-80 g) were induced to lay eggs by injecting with 500 IU per 75 g body weight of human chorionic gonadotrophin (Chorulon) into the dorsal lymph sac, 16 hours before eggs were required. They were placed in a dark incubator overnight in pairs in small holding tanks containing filtered tap water. After egg laying commenced, each female was placed in 1 L of 1 x Marc’s modified ringer’s solution ((MMR: 100 mM NaCl, 2 mM KCl, 1 mM MgSO_4_, 2 mM CaCl_2_, 5mM HEPES, 0.1 mM EDTA pH 8.0) to prevent activation. Eggs were collected hourly and fertilised using fresh testes from a euthanised adult *X. laevis* male. Egg jelly was removed immediately following embryo rotation (15-20 minutes), using a 2% solution of L-Cysteine (pH 7.9) and the resulting de-jellied embryos were rinsed three times in MMR. Embryos were raised in small batches in 0.1 x MMR in 10 cm diameter petri dishes with no antibiotics at 18 ºC and staged according to Nieuwkoop and Faber’s staging series (Nieuwkoop & Faber 1994). Normal development of *X. laevis* embryos and tadpoles was also according to (Nieuwkoop & Faber 1994). All procedures were approved by the University of Otago’s Animal ethics committee under AUP 19/01.

### 2.2 Preparation and delivery of the seizure-inducing chemicals pentylenetetrazole (PTZ) and 4-aminopyridine (4-AP), and the anti-seizure chemical valproic acid (VPA)

All chemicals were freshly prepared before every scheduled experiment. The concentration of these chemicals used in this study was based on previously published information for *Xenopus* tadpoles (Hewapathirane *et al*. 2008). PTZ (Sigma Aldrich, cat# P6500) was prepared as a 100 mM stock solution. Individual tadpoles were placed in 1.2 ml of 5 mM PTZ in 0.1x MMR. Similarly, 5 mM and 10 mM of VPA (sodium salt, Sigma Aldrich, cat# P4543) were prepared from 100 mM stock solution. 0.5 mM of 4-AP (Sigma Aldrich, cat#275875) was prepared from 10 mM stock solution.

### 2.3 Recording of stage 47 tadpole behaviour in 24-well plates

Individual tadpoles were transferred to each well of a 24-well culture plate (Cellstar, Greiner bio-one) (Fig. 1) using a cut down, wide bore glass pipette. The wells were filled with 1.2 ml of 0.1 x MMR, with or without chemicals, so that the tadpoles were submerged. The tadpoles were recorded at 20 ºC using a Panasonic DMC-FZ1000 camera set in movie mode at 25 frames per second. Frames were captured at 1280x 720 pixels at 10 Mbps. Plates were illuminated from below using white LED light source (Neewer 176 ultra bright ϕ5 LED, K5600) and a white perspex diffuser, and the camera was fixed above the well plate with a tripod (Figure 1). Where stated, PTZ, 4-AP and/or VPA were added to wells just prior to the start of video recording. Thirty minutes of video was captured in MP4 format and converted into MPEG2 format for the analysis using Adobe Media Encoder CC 2019 software. All recordings were done between 8am and 6pm with controlled lighting.

**Figure 1:**
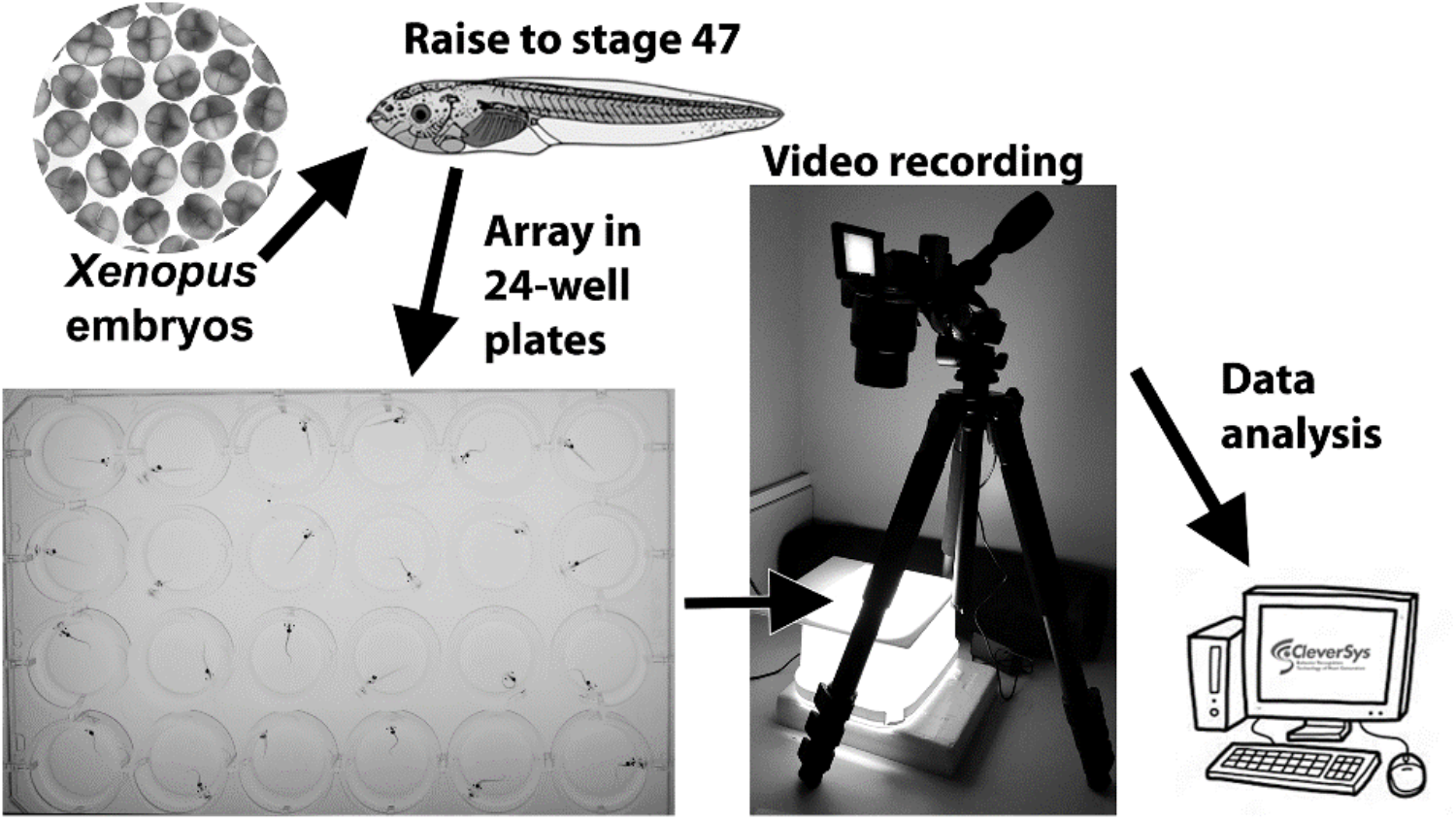
Pipeline for behavioural analysis of tadpoles induced to acute seizure activity with PTZ or 4-AP. *X. laevis* wild type embryos were raised to stage 47, at which point they will progress no further without feeding. Each tadpole was placed in its own well in a 24-well plate and behaviour recorded for 30 minutes. Videos were analysed offline using TopScan software.

### 2.4 Analysis of videos using TopScan (CleverSys)

To analyse seizure behaviour in tadpoles, the TopScan-TopView behaviour analysing system software (CleverSys Inc., USA) was used. Modules *Locoscan, DrugAbuse* and *Circling* were used to quantify seizure-related behaviour in tadpoles. In TopScan, the images of tadpoles were subtracted to create a clear background with just the 24-well plate. Arenas were defined as individual tadpole wells, and were created by using *Arena Design* toolbar. This enables simultaneous analysis of each individual tadpole’s behaviour. For quantification of distance travelled and velocity, the well diameter, which is 17 mm, was entered as 17 cm, and parameters based on distance travelled were corrected *post hoc* by dividing by 10. Event rules and output parameters were then set up to detect and quantify the behaviours of tadpoles such as darting, circling, velocity and movement trajectories to enable phenotyping. A step-by-step procedure of using TopScan software to generate behavioural data from recorded video can be found in supplementary file 1.

### 2.5 Study design

This study was not pre-registered. Tadpoles from a sibship were arbitrarily assigned to treatment groups by pipette transfer. No blinding was performed. To establish seizure-related behaviour in *Xenopus* tadpoles, two seizure-inducing chemicals (PTZ and 4-AP) and one anti-seizure chemical (VPA) were included in this study. Untreated and a VPA-only treated control group were also included. Tadpole treatment groups included in this study are as follows: Untreated, 5 mM VPA, 5 mM PTZ, 5 mM PTZ and 5 mM VPA, 0.5 mM 4-AP, 0.5 mM 4-AP and 5 mM VPA, and 0.5 mM 4-AP and 10 mM VPA. All tadpoles were at stage 47 at the time of analysis, and data for each group was merged from multiple sibships and videos. No sample calculation was used to predetermine sample size, the study was exploratory, and only normally developing tadpoles were included, with exclusion of <1% for any group.

### 2.6 Data analysis

Ethograms were constructed from manually curated data from 10 tadpoles for each group using a Fruchterman–Reingold algorithm and generated using qgraph package v1.9.2 (Epskamp *et al*. 2012). Automatically generated behavioural data from different tadpole sibships was merged for automated analysis. Graphs and statistical analyses were calculated using Prism v9.1.2. D’Agostino-Pearson normality tests were performed on all data (Supplementary data file 2). Comparison between two treatment group means was performed using unpaired t-tests for normally distributed data or Mann-Whitney tests for non-normal data. Comparisons for three or more groups with normal distribution used 1-way ANOVA with Tukey’s post hoc multiple comparison test for all means or a Kruskal-Wallis test with Dunn’s multiple comparisons for non-normal data. Data were presented as mean ± standard error of the mean (SEM) and the statistical significance threshold was set to p < 0.05.

## 3.0 Results

### 3.1 Distinct behaviours are elicited by the acute seizure inducing chemicals PTZ and 4-AP

In rodent models, different seizure behaviours are elicited with different seizure-inducing chemicals. PTZ is associated with tonic-clonic seizures and status epilepticus, and functions via blockade of GABA-A receptors (Loscher 2011). 4-AP acts as a potassium channel blocker, which causes increased neuronal excitability and release of glutamate, leading to status epilepticus (Traynelis & Dingledine 1988; Kobayashi *et al*. 2008). To determine how the tadpole model responds to these drugs, 30 minutes of video data for ten tadpoles that were either untreated or exposed to PTZ or 4-AP were examined manually. Untreated stage 47 tadpoles (controls) were nominally active, usually remaining with the head facing the well side, passive drifting in the well, or swimming slowly for a brief time. This normal tadpole behaviour was termed non-seizure (NS). In contrast, tadpoles treated with the acute seizure-inducing drugs PTZ or 4-AP showed increased levels of activity, manifesting as distinct swimming behaviours which were not seen in untreated tadpoles (Table 1, Fig. 2A). Characteristic “C-shaped contractions” (CSC) were consistently observed, as previously described (Hewapathirane *et al*. 2008). In CSC, the trunk muscles on one side undergo rapid and strong contractions, and as a result the tail bends so far to one side that the tip touches the tadpole’s head (Fig. 2A). However, this was not the dominant or only behaviour elicited: in tadpoles treated with 5 mM PTZ only 7.6% of recorded events were CSC, and for 0.5 mM 4-AP-treated tadpoles it was 12.4% (Table 1). In the remaining time, tadpoles were either behaving normally (NS), or exhibiting one of three distinct behaviours (Fig. 2A). These were termed “uncoordinated tail bends” (UTB), where the tadpole head remained still at the side of the well but the tail was rapidly beating from side to side, “circling” (rapid swimming around the well perimeter), and “darting” (rapid swimming with frequent changes in direction). UTB was the dominant behaviour in 4-AP treated tadpoles, and PTZ treated tadpoles were most likely to be circling (Table 1 and Fig. 2).

**Table 1:** Percentage incidence of each behavioural state ± SEM, for N=10 tadpoles per group over 30 minutes.

|  | Untreated | PTZ Treated | 4-AP Treated |
| --- | --- | --- | --- |
| **Non-seizure (NS)** | 100 | 28.7 ± 2.1 | 13.6 ± 2.2 |
| **Circling** | 0 | 29.0 ± 3.3 | 24.6 ± 2.9 |
| **Darting** | 0 | 22.4 ± 1.8 | 22.7 ± 4.2 |
| **Uncoordinated tail bends (UTB)** | 0 | 12.3 ± 2.8 | 26.8 ± 2.4 |
| **C-shaped contractions (CSC)** | 0 | 7.6 ± 2.3 | 12.4 ± 2.5 |

**Figure 2:**
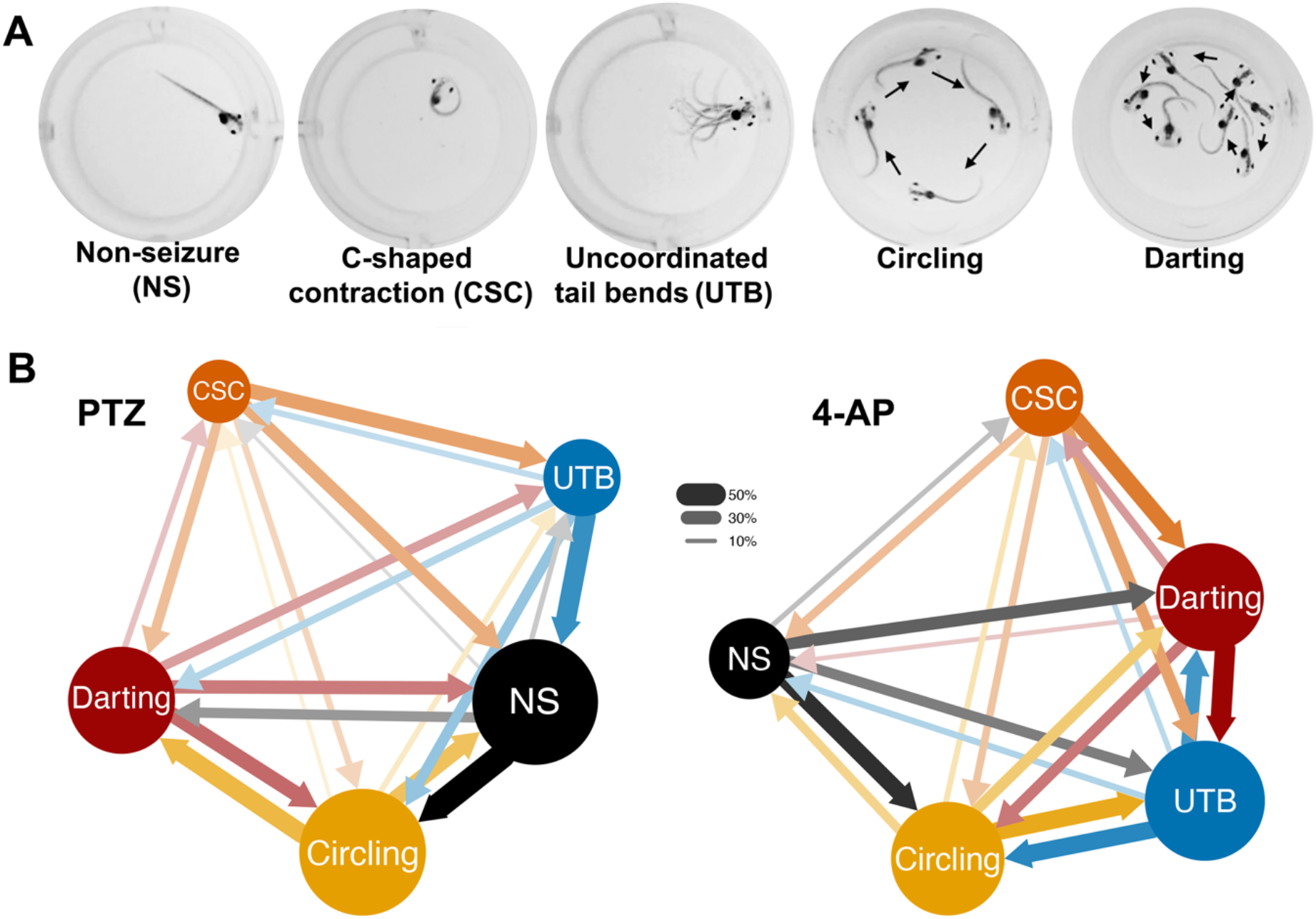
PTZ and 4-AP induce different patterns of swimming behaviour. (A) Typical behaviours of PTZ- and 4-AP-treated tadpoles. Multiple consecutive frames are overlayed for UTB, Circling and Darting, with arrows indicating the sequence of frames. (B) Ethograms created by manual analysis of 10 tadpoles treated with either 5 mM PTZ or 0.5 mM 4-AP to induce acute seizures. Circle diameter represents the mean frequency of each behaviour (see Table 1). Transitions between different behaviours are represented by arrows. Thicker arrows and closer circles indicates transitions between the two behaviours are more frequent. Data can be found in Table 1 and the supplemental file 2.

The frequency of transitions between one behaviour type and another was also recorded and used to construct an ethogram (Fig. 2B). 4-AP-treated tadpoles were most active, and also spent more time in CSC and UTB behaviours than PTZ-treated tadpoles. In PTZ-treated tadpoles, immobile state was followed by circling behaviour 57% of the time, and circling was most often followed by darting (42%) UTB were followed by non-seizure behaviour 45% of the time and by CSC 14% of the time. (Fig. 2C). 4-AP-treated tadpoles did not follow the same pattern as PTZ-treated tadpoles, with darting and circling behaviour frequently alternating with UTB.

### 3.2 Automated behavioural analysis of video recordings with TopScan can be used to track movement and to detect circling and darting behaviour

Manual counting of behaviours from video recordings, while informative, is time-consuming and not scalable to high throughput applications like drug testing. To determine if we could automate this process for drug testing, a pilot study with N=10 tadpoles was conducted. As we could not reliably detect CSC or UTB with TopScan, we compared manually counted CSC to automated analysis of tadpole swimming using TopScan software (Fig. 3). PTZ and 4-AP have been previously shown to induce CSC in stage 47 *Xenopus* tadpoles (Hewapathirane, 2008). Manually counted data confirmed that PTZ-induced tadpoles had an average of 13.4±5.2 CSC during the 30-min recording period, compared to 9.6±2.8 for the cohort treated with both 5 mM VPA and PTZ (Fig. 3A). 4-AP-treated tadpoles had 43.3±7.8 CSC and this was significantly more than groups with 4-AP plus either 5mM VPA (8.4±8.0) or 10mM VPA (7.0±5.70). Both VPA treatment groups had <20% of the CSC seen in PTZ only controls (Fig. 3B).

**Figure 3:**
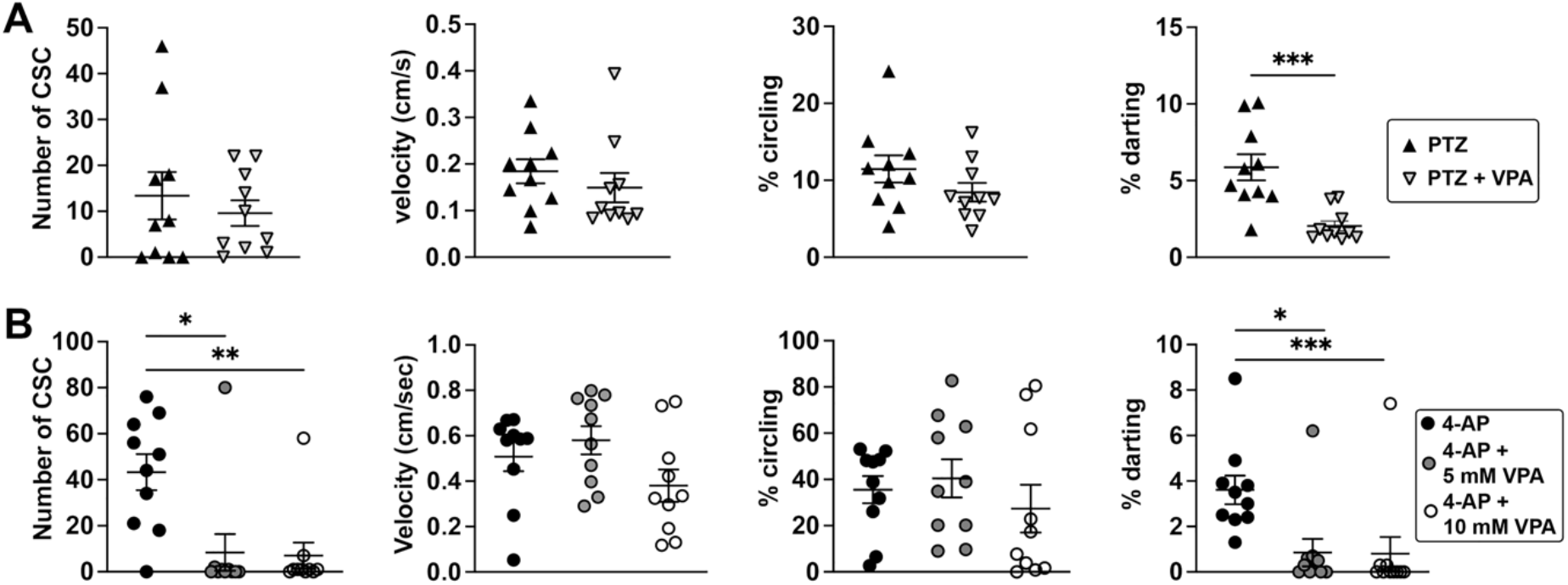
The anti-seizure drug VPA reduces 4-AP-induced C-shaped contractions (CSC) and 4-AP- and PTZ-induced darting behaviour. Scatterplots showing manually counted CSC, mean velocity of tadpoles (cm/s), percentage of time spent circling and percentage of time spent darting. Black line indicates the mean and error bars are SEM. (A) Tadpoles treated with 5 mM PTZ with or without 5 mM VPA, N=10 for each group, compared using unpaired t-test (CSC, circling, darting) or Mann-Whitney test (velocity). (B) Tadpoles treated with 4-AP, with or without VPA at two concentrations, 5 and 10 mM. Statistical analysis was by 1-way ANOVA with Tukey’s post hoc test of all means (circling) and by Kruskal-Wallis with Dunn’s post hoc test (CSC, darting, velocity). N=10 tadpoles in each group. Statistically different groups are indicated by asterisks: * p<0.05, ** p<0.01, *** p<0.001. Data can be found in the supplemental file 2.

The same videos were also analysed with TopScan to determine mean swimming velocity (cm/s) and the percentage of time spent circling or darting. Of the automatically scored behaviours, VPA only resulted in a significant reduction in the percentage of time tadpoles spent darting (Fig. 3A,B).

### 3.3 High-throughput automated analysis confirms the efficacy of VPA in reducing seizure-related activity

Having shown the potential of automated behavioural analysis to quantify the effect of seizure-inducing chemicals PTZ and 4-AP, at least for darting, we increased the number of tadpoles in the analysis. We also included untreated tadpoles (controls) and tadpoles treated with 5mM VPA in this analysis (Fig. 4). As CSC could not be reliably detected by TopScan, we sought to use the other abnormal behaviours to assess the effectiveness of anti-seizure drugs, using VPA as a benchmark. Movement trajectories (tracks) of individual tadpoles confirmed the increased activity of PTZ and 4-AP treated tadpoles (Fig. 4A). Further, the co-administration of VPA reduced the excessive movement of the tadpoles to control levels. Notably, 4-AP-treated tadpoles were still more active than controls even in the presence of 10 mM VPA (Fig. 4A).

**Figure 4:**
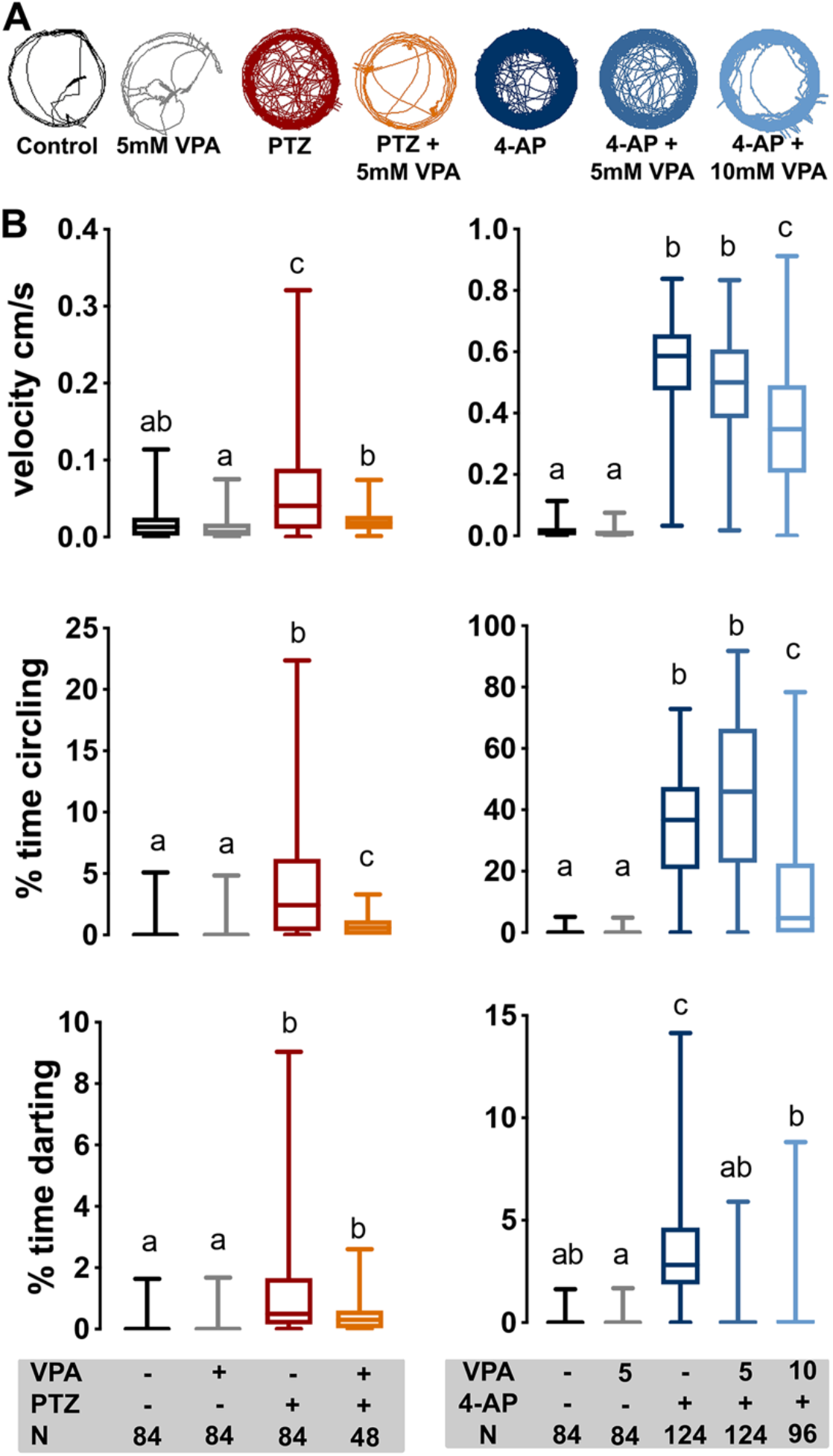
Automatic analysis shows that VPA reduces PTZ- and 4-AP-induced swimming behaviours. A) Representative examples of individual tadpole movement trajectories each treatment group, over the 30 minutes of the trial (treatment is colour-coded as in B). B) Box plots of average tadpole swimming velocity (cm/sec), percentage of time spent swimming in rapid circles and percentage of time spent darting. Middle line indicates the median, upper and lower lines indicate the upper and lower quartile, and the whiskers indicate the minimum and maximum values. Kruskal-Wallis test with Dunn’s post hoc testing of all means was used to analyse each dataset. Groups that share a lower case letter are not significantly different from one another whereas groups which are significantly different are indicated by a different lower case letter. N=number of tadpoles, VPA was used at 5 or 10 mM, PTZ at 5 mM and 4-AP at 0.5 mM). Raw data can be found in supplemental file 2.

All three measured parameters: mean velocity, percentage of time circling and percentage of time darting, were significantly increased in either 5 mM PTZ- or 0.5 mM 4-AP-treated tadpoles (Fig. 4B). Circling and darting behaviour was rarely seen in control treatment groups (< 0.2% and <0.1% respectively). PTZ-induced behaviours were all significantly lower in the cohort with co-administered 5 mM VPA. Mean velocity was 3.3-fold lower, the percentage of time spent circling was 5.7-fold lower, and the percentage of time darting was 2.9-fold lower in the presence of VPA (Fig. 4B).

Rapid swimming induced by 4-AP, seen as velocity and circling measures, gave mixed results with 5 mM VPA, but tadpoles co-treated with 4-AP and the higher dose of 10 mM showed a significant reduction in both behaviours (2.2-fold less time circling, 1.5-fold lower mean velocity). 4-AP-induced darting was not seen when VPA was present at either concentration (Fig. 4B).

### 3.4 VPA treatment delays the time to first circling or darting event induced by 4-AP, but only delays the onset of circling induced by PTZ

To test whether VPA affects the onset time of PTZ- and 4-AP-induced circling and darting behaviour, we recorded the time of the first darting or circling event for each tadpole in the study and presented this as a survival plot (Fig. 5). Tadpoles that did not have any events in the 30 minute recording window were censored and the percentage for each group can be found in Table 2. An increased time to the first seizure event in the presence of an ASM may indicate elevated seizure threshold. In PTZ-treated tadpoles, the latency to first circling event was significantly higher in the presence of VPA and the percentage of tadpoles that never circled more than doubled, from 13 to 29% (Fig 5A). All 4-AP induced tadpoles had at least one circling event, whereas 7 or 27% of tadpoles co-treated with 5 or 10mM VPA respectively had no events in the 30 minute period (Fig 5B). PTZ induced darting behaviour was not affected by VPA (Fig.5C). 4-AP induced darting was significantly delayed by either dose of VPA, with only 12% of 4-AP tadpoles having no darting compared to 60-61% of tadpoles co-treated with 5 or 10 mM VPA (Fig. 5D).

**Table 2:** Percentage of tadpoles showing no circling or darting behaviour for each treatment group

| Treatment group | Sample size | % with no circling | % with no darting |
| --- | --- | --- | --- |
| PTZ | 84 | 13.1 | 15.5 |
| PTZ and VPA | 48 | 29.2 | 20.8 |
| 4-AP | 124 | 0.0 | 12.1 |
| 4-AP and 5mM VPA | 124 | 6.5 | 61.3 |
| 4-AP and 10mM VPA | 96 | 27.1 | 60.4 |

**Figure 5:**
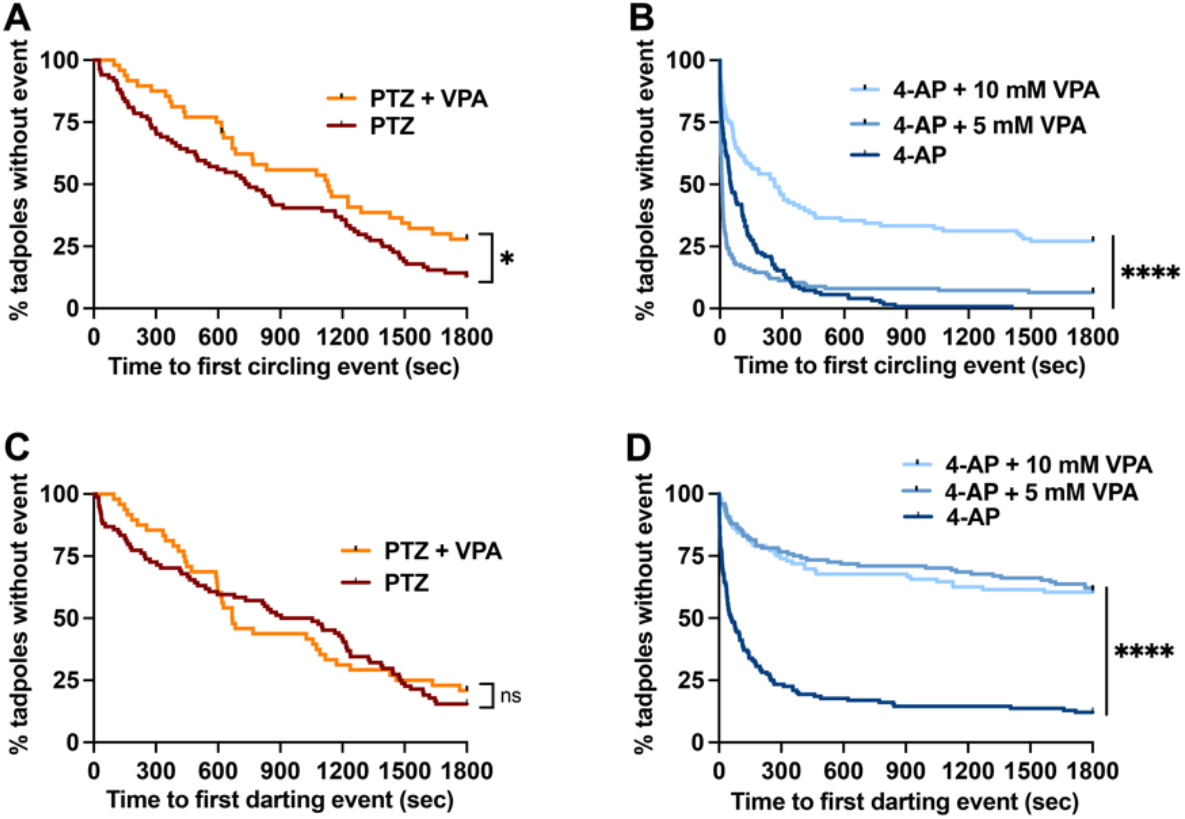
Time to first 4-AP induced circling or darting event is longer in the presence of anti-seizure drug VPA. Kaplan-Meier plots where time to first seizure event (circling or darting) has been substituted for death. Tadpoles were censored if they did not have any events in the 30 minute timespan and curves were compared with log-rank (Mantel-Cox) test. A) Time to first circling event was longer for PTZ tadpoles co-treated with VPA. (χ^2^ = 4.5, df= 1, p=0.033) B) Tadpoles induced with 4-AP all had at least one circling event, but VPA increased latency to first event, and this effect increased with dose (χ^2^ =63.3, df=2, p<0.0001). C) The time to first darting effect induced by PTZ was not altered by VPA (χ^2^ = 0.1, df=1, p=0.740) D) Time to first darting event induced by 4-AP was significantly longer in the presence of VPA (χ^2^ = 129.2, df=2, p<0.0001) Group sizes: PTZ N=84, PTZ + VPA N=48, 4-AP N=124, 4-AP + 5mM VPA N=124 4-AP + 10 mM VPA, N=96. Raw data can be found in supplemental file 2. * p<0.05, **** p<0.0001.

## 4.0 Discussion

### 4.1 PTZ and 4-AP treated tadpoles elicit similar patterns of seizure-related behaviours

PTZ and 4-AP are widely used chemicals to induce acute seizures in various mammalian and non-mammalian animal model systems (Hewapathirane *et al*. 2008; Ellis *et al*. 2012; Cunliffe *et al*. 2015; Johan Arief *et al*. 2018). Depending upon the animal model system and concentration, PTZ and 4-AP induce seizures within few minutes and the epileptic effect can continue for hours. Studies have shown that both latency to seizure and seizure severity after PTZ treatment is dose-dependent (Klioueva *et al*. 2001; Hewapathirane *et al*. 2008; Luttjohann *et al*. 2009; Pineda *et al*. 2011; Van Erum *et al*. 2020). PTZ induces acute seizures by antagonising GABA-A receptors, which reduces inhibitory GABA signalling (Loscher 2011). 4-AP acts via a different mechanism, blocking potassium channels (Traynelis & Dingledine 1988; Kobayashi *et al*. 2008), and is also proven to induce spontaneous seizure-like events in *X. laevis* and zebrafish larval models (Hewapathirane *et al*. 2008) (Winter *et al*. 2017).

While PTZ and 4-AP induced the same set of stereotypic behaviours, our ethogram revealed inducer-dependent distinct sequences of behavioural events (Fig. 2B). C-shaped contractions were detected for both groups, but could only be manually scored due to limitations with TopScan (primarily the low frame rate). CSC likely arise as a result of neuronal excitation spreading to the networks of the tadpole brain normally dedicated to the well characterised C-start kinetic response (Karin v & Wassersug 2000). The other stereotypical behaviours we describe likely also arise from similar overstimulation of inherent motor pattern control networks in the tadpole brain. Hence, these behaviours are not unique to acute seizure models and we are detecting them due to their exaggerated or prolonged manifestations. In zebrafish larvae, a functional study comparing several acute seizure inducing chemicals showed that PTZ resulted in neural activity that was strongest in the rhomencephalon (hindbrain) and 4-AP the tectum (Winter *et al*. 2017).

### 4.1 Automated detection of seizure behaviours in tadpoles shows that darting is the most sensitive indicator of drug efficacy

The primary clinical diagnostic test for confirming epileptic seizures in humans is via analysis of electroencephalogram (EEG) data. However, the occurrence of abnormal behavioural movements is generally the first indicator of epilepsy and is an integral part of diagnosis (Fisher *et al*. 2017; Falco-Walter *et al*. 2018). In a pioneering study, Hewapathirane et al (2008) described a series of progressive seizure-related behaviours (class I to V) using manual video analysis of *Xenopus laevis* tadpoles induced with epileptogenic drugs (Hewapathirane *et al*. 2008). Under normal conditions, tadpoles show slow swimming and brief pause-like behaviour which we also observed here, in both untreated and VPA only treated control groups of tadpoles. Tadpoles tend to display abnormal swimming behaviour such as increased locomotor activity/agitation (also termed as hyperlocomotion) as a sign or pattern of seizure progression. Circling and darting (as described and detected here) best represent class I behaviour, defined as ‘*bouts of intermittent rapid swimming, involving swimming in tight circles or sporadic rapid swimming with abrupt changes in direction (“darting”)’*. However there is also some overlap with class II and class III behaviours (class II: as class I but with intermittent immobility, class III: additional loss of postural control, e.g. tadpole is on its side or upside down). In contrast, UTB are most like class IV, and CSC are class V, representing the culmination of a generalised seizure (Hewapathirane *et al*. 2008). Therefore we have been able to automate detection of previously described class I-III behaviours, using “circling” and “darting” but not class IV or V. Time spent in both circling and darting behaviours was significantly higher in tadpoles treated with PTZ and 4-AP compared to control tadpoles, and treatment with VPA significantly reduced the number of circling and darting bouts (Fig. 4). We also used mean velocity as a measure of seizure-induced hyperactivity. This has not previously been used as a measure for seizure activity in tadpoles, but hyperlocomotion has been described in zebrafish larvae (Baraban *et al*. 2005; Afrikanova *et al*. 2013; Gawel *et al*. 2021). Together, velocity, darting and circling were useful indicators of seizure activity induced by either PTZ or 4-AP, but only when large cohorts were analysed. Automation of behaviours should therefore be useful in analysing the effects of variants of unknown significance in epilepsy causing genes.

Time spent engaged in darting behaviour was the most sensitive method of detecting reductions in acute seizure behaviours by the anti-seizure drug VPA in a small cohort (Fig. 3). The broad anti-epileptic effect of VPA is due to a combination of effects that act to both increase inhibitory GABA signalling and decrease excitatory signals such as glutamate (Romoli *et al*. 2019). The latency, severity, nature and progression of seizure-related behaviour seen in this study is similar to the previous published studies in *Xenopus* tadpoles, zebrafish and rodents. For example, tonic/clonic and uncontrolled motor movements seen in mice (Van Erum *et al*. 2020), rats (Luttjohann *et al*. 2009), zebrafish (Afrikanova *et al*. 2013; Gawel *et al*. 2021) and other models (Johan Arief *et al*. 2018) after PTZ can be correlated to increased velocity as well as time spent in darting and circling behaviour seen in this study.

### 4.2 *Xenopus* tadpole as a model organism for epilepsy, clinical significance and future directions

Several studies have been conducted in the past for the purpose of using zebrafish (*Danio rerio*), nematode worms (*Caenorhabditis elegans*) or fruit flies (*Drosophila melanogaster*) as a potential non-mammalian models for seizure or seizure-like activity (Hortopan *et al*. 2010; Cunliffe *et al*. 2015; Johan Arief *et al*. 2018; Afrikanova *et al*. 2013; Zhu *et al*. 2020; Gawel *et al*. 2021). The main idea is to establish a model which is less time-consuming, less expensive, robust and straight forward for high-throughput anti-epileptic drug discovery. With the exception of zebrafish, these studies quantified behavioural seizures or seizure progression just in terms of locomotion/movement, as we have done here. While this is a good first step, if non-mammalian models are to be useful for epilepsy research, seizure-related behaviours will need to be validated by identifying neural signatures of seizures (e.g., electrophysiological brain recordings (Hewapathirane *et al*. 2008) or seizure-induced changes in gene expression (Labiner *et al*. 1993; Ribak *et al*. 1997). In both zebrafish larvae, and *X. laevis* tadpoles, electrophysiological recording from the larval brain has been demonstrated (Diaz Verdugo *et al*. 2019; Baraban *et al*. 2013; Hewapathirane *et al*. 2008). Furthermore, this activity has been expressly linked to calcium signalling in both models (Hewapathirane *et al*. 2008; Diaz Verdugo *et al*. 2019). As per our current understanding, this is the first study to automatically quantify seizure-related behaviour in *Xenopus* tadpoles and should be useful to unlock the potential of *X. laevis* tadpoles as models of genetic epilepsy.

## 5.0 Concluding remarks

We quantified behaviours elicited in *X. laevis* tadpoles by the acute seizure-inducing chemicals PTZ and 4-AP and showed that these behaviours were eliminated or reduced by the anti-seizure drug VPA. To develop a new rapid pre-clinical *Xenopus* model for chronic epilepsy where a genetic basis is suspected, characterization and quantification of seizure-related behaviour will be crucial, as it provides a basis for triaging variants before undertaking more time-consuming electrophysiological experiments. We suggest that the automated behaviour analysis described here could be a useful first step in the development of much needed pre-clinical non-mammalian models of genetic epilepsy.

## Abbreviations

4-AP: 4-aminopyridine
ASM: anti-seizure medication
CRISPR: Clustered regularly interspaced palindromic repeats
CSC: C-shaped contractions
DEE: developmental and epileptic encephalopathy
EEG: electroencephalogram
LED: light emitting diode
MMR: Marc’s modified ringers
NS: non-seizure behaviour
PTZ: pentylenetetrazole
SEM: standard error of the mean
UTB: uncontrolled tail bends
VPA: valproic acid.

## Data Availability Statement

A detailed methodology and a spreadsheet of all raw data and statistical testing results can be found in the supplementary information.

Supplementary file 1: detailed TopScan protocol and behavioural parameters

Supplementary file 2: Raw data and statistical analysis supporting the conclusions

## Ethics Statement

All experiments were performed in accordance with ARRIVE guidelines. *Xenopus* embryos were produced using protocol approved by the University of Otago Animal Ethics Committee under AUP 19/01 and tadpoles killed by immersion in overdose of tricaine. No measures were taken to minimize pain or discomfort in tadpoles or adults, besides gentle handling.

## Author Contributions

Conceptualization, methodology, investigation, SP, formal analysis, data curation: SP, CB, PC; Writing: SP, PS, CB; Supervision: PS and CB; Project administration and funding acquisition: CB.

## Acknowledgments

The authors would like to thank Nikita Woodhead for care of the *Xenopus* colony, and Joanna Ward for laboratory support. We also thank Cabriana Earl for critical reading of the manuscript, and Sulagna Banerjee for assistance with setting up TopScan for *Xenopus* tadpoles.

